# EEG spectral attractors identify a geometric core of resting brain activity

**DOI:** 10.1101/2023.10.13.562264

**Authors:** Parham Pourdavood, Michael S. Jacob

## Abstract

Spectral analysis of electroencephalographic (EEG) data simplifies the characterization of periodic band parameters but can obscure underlying dynamics. By contrast, reconstruction of neural activity in state-space preserves geometric complexity in the form of a multidimensional, global attractor. Here we combine these perspectives, inferring complexity and shared dynamics from eigen-time-delay embedding of periodic and aperiodic spectral parameters to yield unique dynamical attractors for each EEG parameter. We find that resting-state alpha and aperiodic attractors show low geometric complexity and shared dynamics with all other frequency bands, what we refer to as geometric cross-parameter coupling. Further, the geometric signatures of alpha and aperiodic attractors dominate spectral dynamics, identifying a geometric core of brain activity. Non-core attractors demonstrate higher complexity but retain traces of this low-dimensional signal, supporting a hypothesis that frequency specific information differentiates out of an integrative, dynamic core. Older adults show lower geometric complexity but greater geometric coupling, resulting from dedifferentiation of gamma band activity. The form and content of resting-state thoughts were further associated with the complexity of core dynamics. Thus, the hallmarks of resting-state EEG in the frequency domain, the alpha peak and the aperiodic backbone, reflect a dynamic, geometric core of resting-state brain activity. This evidence for a geometric core in EEG complements evidence for a regionally defined dynamic core from fMRI-based neuroimaging, further supporting the utility of geometric approaches to the analysis of neural data.

## Introduction

Since their discovery, electrophysiologic signals have been defined by rhythmicity. This has led to a “rhythms of the brain” approach to studying physiologic processes, by linking putative cognitive functions to indices of frequency-based activity, such as power, phase coherence, etc. (Buzsaki, 2006). This approach has been expanded by growing evidence that interactions between frequency bands, so-called cross-frequency coupling, underlie cognitive function (Aru et al., 2015; Canolty & Knight, 2010; Hyafil et al., 2015; Palva & Palva, 2018) and that disruptions in synchronized activity contribute to neuropsychiatric dysfunction (Grover et al., 2021; Mathalon & Sohal, 2015). Cross-frequency interactions are characterized by pairwise measures of coherence and/or correlation, often interpreted as connectivity since they are thought to reflect functional interactions between local or global brain networks, depending on the frequency bands involved (Florin & Baillet, 2015; Marzetti et al., 2019). Analytic pitfalls notwithstanding (Hyafil, 2015), these approaches generally emphasize linear assessment of time-frequency decomposition linked to a stimulus or task (Cao et al., 2022; Hülsemann et al., 2019). Emerging evidence for the importance of aperiodic activity (not defined by a distinct frequency) complicates analysis of cross-frequency interactions, (Canales-Johnson et al., 2021; Donoghue, Dominguez, et al., 2020; Donoghue, Haller, et al., 2020; Schaworonkow & Voytek, 2021; Tröndle et al., 2023) raising the necessity of identifying new methods to investigate interactions between aperiodic and periodic parameters. Moreover, time-frequency coupling approaches are difficult to apply to resting-state paradigms, since intrinsic coupling is not necessarily linked to an experimental variable (Engel et al., 2013).

In parallel with these developments in the study of brain rhythms, alternative perspectives have examined brain activity as a multidimensional signal, with complex, nonlinear and nonstationary dynamics (Breakspear, 2017; Stringer et al., 2019; Tozzi, 2019). Although signal complexity of this nature is often dismissed as random variance, it may contribute meaningfully to cognition and behavior as part of a globally integrated, dynamical system (Busch et al., 2009; Galgali et al., 2023; Waschke et al., 2021). One approach to analytically utilize this dynamical systems perspective is to embed physiologic signals onto a multidimensional, state-space manifold (Freeman, 1987; McKenna et al., 1994; Tozzi & Peters, 2017; Tsuda, 2001). When a system’s states converge around a low-dimensional set of states, this suggests that the system may be operating in an attractor basin (Breakspear, 2017). These methods have ushered in a ‘geometric approach’ to analyzing the dynamic signatures and motifs over activity space (Azeredo da Silveira & Rieke, 2021; Chung & Abbott, 2021; Kriegeskorte & Kievit, 2013). Although state-space embedding procedures have long been used to investigate EEG activity (Freeman, 1987; Tsuda, 2001), such approaches have not been employed to investigate cross-parameter coupling. Manifold-to-manifold or attractor-to-attractor mapping relationships might provide a novel approach to investigating the structure of EEG, the interactions between EEG parameters and especially where traditional cross-frequency coupling approaches may not be appropriate.

Geometric approaches have helped identify regional dynamics from fMRI that are common across a range of cognitive tasks (Shine et al., 2019). These shared dynamics cut across traditional functional networks and are based instead on unique, topological signatures in the dynamics of activity. The resulting regions and their dynamical manifolds have been reported as evidence of an integrative “dynamic core” in the brain. The concept of an integrative core is originally attributed to Varela (Varela, 1995), who proposed that specific cognitive operations reflect transient synchronization that emerges from ongoing activity in a dynamically unstable dominant assembly (Le Van Quyen, 2003; Varela, 1995). The term “dynamic core” was later developed by Tononi and Edelman to reflect a continually varying, unified process that yields high degrees of differentiation and complexity (Tononi & Edelman, 1998). Given the promise of geometric, manifold based representations of neural data, we reasoned that the dynamic core of EEG might be evident in shared dynamics across the frequency domain, as an alternative to traditional frequency-based synchronization approaches. To define the geometry and ascertain the complexity of EEG spectral dynamics, we used state-space embedding procedures and a measure of geometric cross-parameter coupling during a resting-state paradigm. We find that the dynamics of the alpha band and the aperiodic exponent contain a low-dimensional, core-set of dynamically unstable, geometric signatures that predict the dynamics of other EEG parameters. We further examined differences in this geometric core in older adults (OA), and associations with the form and content of resting-state thoughts. Our findings emphasize the inherently unstable nature of the EEG’s dynamic core, whereby higher-frequency non-core operations differentiate out of core-dynamics to yield structures of higher complexity. This work has implications for dynamical systems theories of neural function and models of aging.

## Methods

All of the MATLAB code for the algorithm and analyses of this manuscript is publicly available online in the Github repository named Eigen Manifold Cross Mapping (EMCM; https://github.com/ParhamP/EMCM).

### Dataset

We utilized public access data from the “Leipzig Study for Mind-Body-Emotion Interactions” (LEMON) dataset (Babayan et al., 2019) that includes 227 healthy participants, including young adult participants (YA, N=153, mean= 25.1 years, median=24 years, standard deviation=3.1, 45 females), and older adult participants (OA, N=74, mean= 67.6 years, median=67 years, standard deviation=4.7, 37 females). Participants in this study underwent a two-day assessment in Leipzig, Germany between the years 2013 and 2015. The study included resting EEG and fMRI and completed a battery of neuropsychological and cognitive tests. Participants were free of psychiatric or neurological disorders, for a full list of exclusion criteria see (Babayan et al., 2019).

### EEG Acquisition and Preprocessing

Details of the acquisition and preprocessing are available in (Babayan et al., 2019). The resting-state EEG in the LEMON study was recorded from 216 participants and consisted of 16 resting-state blocks, each lasting 60 seconds. Half of the blocks were performed with the participant’s eyes closed (EC) and the other half with their eyes open (EO). The blocks were visually cued and the participants were seated in front of a computer screen and were instructed to remain awake and focus their gaze on a black cross displayed on a white background during the eyes-open blocks. The recording was done using a BrainAmp MR Plus amplifier, 61 scalp electrodes, and 1 electrode below the right eye. The EEG data was captured using a bandpass filter with a range of 0.015 Hz to 1 kHz and the recordings were then converted into a digital format with a sampling rate of 2500 Hz. At the preprocessing, 13 participants were excluded from the analysis due to missing participant demographic information, different sampling rate, and poor data quality. The raw EEG data was then preprocessed by downsampling to 250 Hz and applying a 1-45 Hz bandpass filter. Outlier channels or data intervals were identified by visual inspection and removed. ICA was utilized to reject eye movements, eye blinks, and heartbeats. Here, we utilized the resting-state EEG data from 203 participants (139 YA and 64 OA).

### EEG Processing

Preprocessed resting-state electroencephalogram (EEG) data for eyes closed (EC) and eyes open (EO) conditions, 61 scalp electrodes (Fp1, Fp2, F7, F3, Fz, F4, F8,FC5, FC1, FC2, FC6, T7, C3, Cz, C4, T8, CP5, CP1, CP2, CP6, AFz, P7, P3, Pz, P4, P8, PO9, O1, Oz, O2, PO10, AF7, AF3, AF4, AF8, F5, F1, F2, F6, FT7,FC3, FC4, FT8, C5, C1, C2, C6, TP7, CP3, CPz, CP4, TP8, P5, P1, P2, P6, PO7, PO3, POz, PO4, PO8). The EEG spectra was then parameterized into canonical periodic oscillations and aperiodic components using the FOOOF (Fitting Oscillations & One Over F) algorithm (Donoghue, Haller, et al., 2020) (Figure 1, B), within a frequency range of 1-45 Hz. We utilized the default parameters for FOOOF, with peak width limits between 1 and 6, peak threshold of 2.0 and ‘fixed’ background mode, which was applied on sliding 2-second epochs of EEG time-series data (one participant containing different sampling rate was omitted). The FOOOF algorithm provided good fits for all participants and TR epochs, with mean *R*^2^ of 0.85 (range 0.76-0.89) for YA and 0.82 (range 0.72-0.89) for OA. This yields aperiodic parameters (slope and intercept) and residual periodic power for each epoch (Figure 1, B). For five participants, there was a missing epoch in the eyes closed condition ([sub-010028, channel C6, TR=503], [sub-010092, Channel C1, TR=554], [sub-010152, Channel PO4, TR=549], [sub-010199, Channel PO9, TR=212], [sub-010255, Channel=FC6, TR=859]). Since none of those channels are part of the set of channels we utilized for our parameter averages, they did not affect our analysis. We extracted EEG periodic power within the following frequency ranges: delta (1-3 Hz), theta (4-7 Hz), alpha (8-12 Hz), beta (15-25 Hz) and low gamma (30-45 Hz), based on the International Federation of Clinical Neurophysiology (Babiloni et al., 2020). For each of the canonical EEG rhythms, an average across a set of channels that contained the highest power was selected for further analysis ({Fz, FC1, FC2, F1, F2, AFz} for delta, theta and gamma, and {Pz, POz, P1, P2, CP1, CP2, CPz} for alpha and beta). For the aperiodic exponent, an average across a set of channels that contained the shallowest slope were selected for further analysis (F1, F3, FC1, FC3, F2, F4, FC2, FC4).

**Figure 1.**
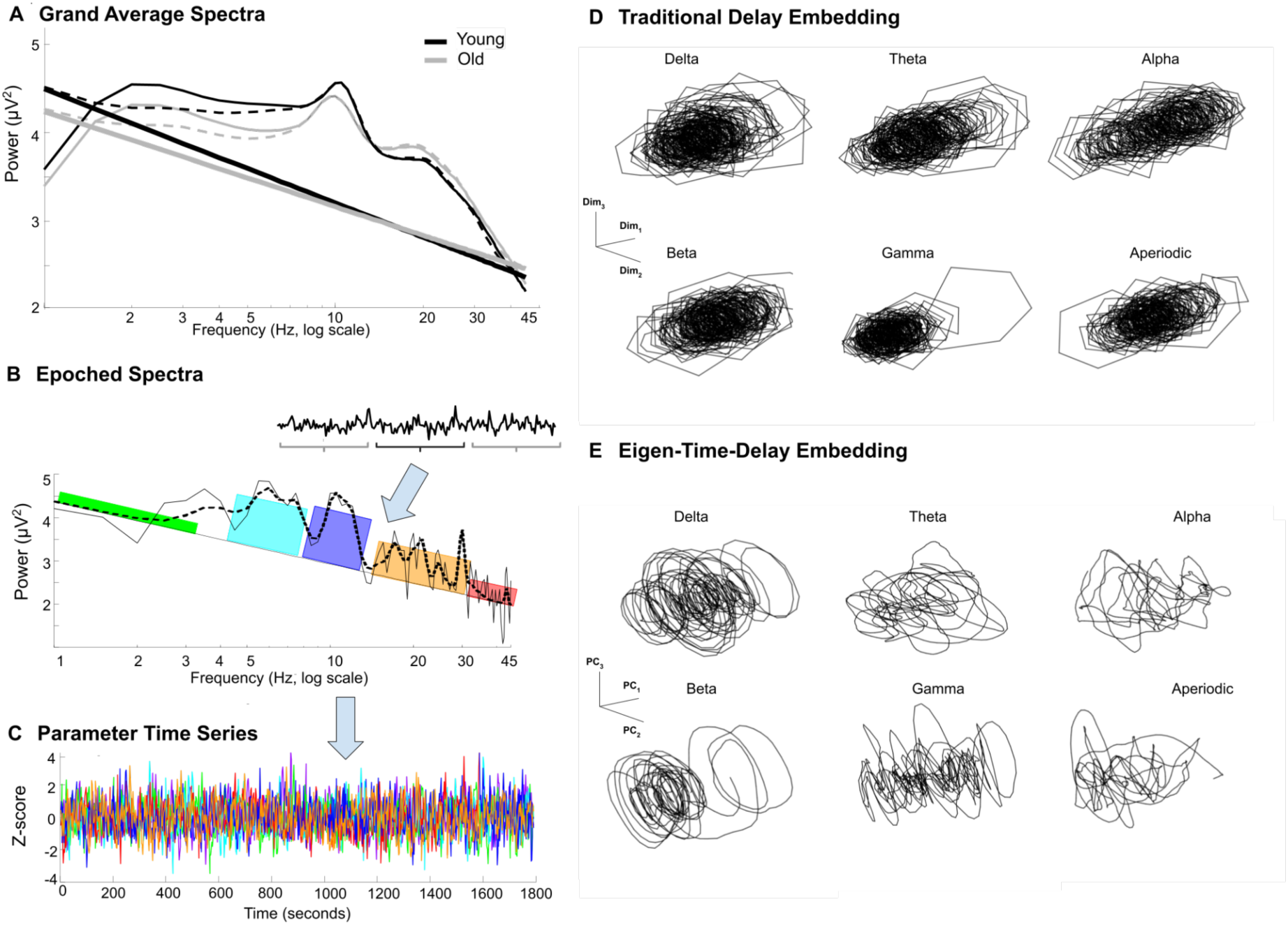
EEG parameterization and embedding methods (A) Grand average spectra during EC for YA (n=138, black) and OA (n=63, gray) participants. Dashed line indicate FOOOF fitting algorithm and the solid, linear fit below indicates the slope of the aperiodic fit for each group. (B) An example of extracted EEG parameters using FOOOF for one epoch, from one participant; light green (delta), cyan (theta), blue (alpha), yellow (beta), and red (gamma). (C) Normalized time-series for each parameter in the same colors from the participant in (B). (D) Each parameter’s time-series is used to reconstruct its high-dimensional attractor using traditional time-delay-embedding. (E) Eigen-time-delayed attractors are plotted from the first three principal components (PC1, PC2, PC3).

### Attractor Reconstruction

The overall intuition for our approach combines recent advances in the linear approximation of otherwise nonlinear attractor systems based on Koopman theory (Brunton et al., 2017) with cross mapping methods to infer causal interactions between attractor manifolds (Sugihara et al., 2012). Based on the assumption that an EEG time-series is a projected measurement of higher-dimensional dynamics in the brain (Tozzi, 2019), our approach begins with a nonlinear state-space reconstruction technique called time-delay embedding to embed EEG time-series in higher dimensional space. Briefly, this approach is based on Takens’ theorem that states under certain conditions, a smooth attractor can be reconstructed from its one dimensional measurement such that it is diffeomorphic to the original attractor. Given time-series *x* (*t*), its embedding in E dimensions can be represented as: *X_E_* = (*x* (*t*), *x* (*t* − τ), *x* (*t* − 2τ), …, *x* (*t* − (*E* − 1) τ), where τ determines how much the original time-series is delayed. The selection of E can be determined directly from the data. Following (Tajima et al., 2015), we choose an embedding dimension at which further increases of E will not lead to increases in the Convergent Cross Mapping score (see below). For the EEG data, we found an average E of 60 across all participants, parameters, and channels. Random noise (Gaussian with mean zero and standard deviation of one) was applied to *X_E_*(*t*) by multiplying a square random matrix from the left in order to reduce autocorrelation and contributions of non-stationarity in the delayed time-series (see also, (Tajima et al., 2015)). Other approaches for determining E were considered, such as the false-nearest neighbor algorithm (Kennel et al., 1992), but it’s been shown to be susceptible to large noisy data (C. Rhodes & Morari, 1997).

After finding an appropriate E, we adopt a technique called Eigen-Time-Delay (ETD) embedding (Juang & Pappa, 1985) and (Brunton et al., 2017) which applies Principal Component Analysis (PCA) to *X_E_*(*t*) and orders the eigen-time-delay vectors by the fraction of variance they contribute to the original phase-space, resulting in *V_E_* = (*ν*_1_, *ν*_2_, *ν*_3_, …, *v_E_*), where *X_E_v_i_* = λ*_i_v_i_* for all eigenvectors *ν*_1_, …, *v_E_* and eigenvalues λ_1_, …, λ*_E_*. This approach to data reduction of non-linear methods has been suggested as a way to reduce noise (Hegger et al., 1999) and further construct linear approximations of the system (Brunton et al., 2017), although this was not further explored in this study. Therefore, our application of PCA to delay-embedding yields phase-space axes that are weighted by their contribution to the overall signal as has been utilized by others to identify the dynamical core (Shine et al., 2019).

### Retained Components and Complexity

In accordance with Kaiser criterion, we retained all the principal components with eigenvalues above 1, which are the components that captured at least as much variance as a single dimension of the original pre-ETD attractor (Kaiser, 1960). Hence, our final reconstructed attractors will be denoted as *V_r_*= (*ν*_1_, *ν*_2_, *ν*_3_, …, *v_r_*), where *r* represents the number of principal components that are retained. Additionally, we designate *r* to serve as a measure of geometric complexity since a larger number of retained components corresponds to a higher amount of statistical variability or signatures in the system. This approach is intuitively similar to the mathematical use of the term topological complexity (Moyal & Edelman, 2019).

### LLE Analysis and Mean Frequency of Principal Components

We utilized Maximal Lyapunov Exponent (MLE) analysis on each ETD-reconstructed dimension to characterize instability (chaotic nature) of their dynamics (Ma et al., 2021; Miller, 1964). MATLAB’s standard “lyapunovExponent” function with default parameters was used to measure the LLE for all the retained principal components of ETD-reconstructed attractors (Rosenstein et al., 1993). The default variables are the normalized frequency of 2π, embedding dimension of 2, lag of 1, minimum separation time of 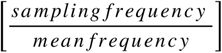, and expansion range of [1, 5]. We calculated the mean frequency of our ETD-reconstructed attractors’ dimensions to characterize their periodicity using MATLAB’s standard “meanfreq” function.

### Geometric Cross-Parameter Coupling

We employ a variant of the Convergent Cross Mapping (CCM) algorithm to assess shared dynamics and causal influence between two dynamical systems (Sugihara et al., 2012). Briefly, CCM uses the k-nearest neighbors (KNN) algorithm in euclidean space to identify the neighbors for all points in each attractor as well as the weight for each neighbor based on euclidean distance. The number of neighbors for each point is equal to *r* + 1, (number of dimensions in the attractor, plus one). For each point in an attractor *V_r_*(*t*), the set of time-indices for its neighbors (arranged in ascending order of distance) and their corresponding weights can be represented as {*t*_1_, *t*_2_, …, *t_r_* _+ 1_} and {*w*_1_, *w*_2_, …, *w_r_* _+ 1_}, respectively, where *w_i_* = *u_i_* / ∑ *u_j_*, for *j* = 1, …, *r* + 1, and *u_i_* = *exp* (− *d* (*V_r_*(*t*), *V_r_*(*i*)) / *d* (*V_r_*(*t*), *V_r_*(*t*_1_)), where *d* is the euclidean distance between the two given points. Subsequently, the algorithm uses the set of neighbors from one attractor to estimate the other and vice versa. For example, we can use attractor 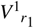’s local neighborhood dynamics to estimate attractor 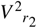, where each point in the estimated attractor can be represented as 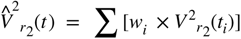, for *i* = 1, …, *r*_1_ + 1. To obtain a score for the degree of shared dynamics and causality, CCM computes the average of all Pearson correlations between the dimensions of the original manifold and its predicted manifold. We use our Eigen modified attractors in CCM to take a weighted average of the correlations across all dimensions based on the eigenvalues from ETD (see the attractor reconstruction section step above). Traditionally, a higher score for the estimation of 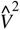 by *V*^1^, is interpreted as *V*^2^ having a causal influence on *V*^1^: if *V*^1^ obtains a precise estimate of *V*^2^, this can be interpreted as *V*^1^ containing local dynamics about *V*^2^, implying *V* ^2^ has previously impacted the dynamics of *V*^1^ by leaving a distinctive geometric imprint (39).

### Statistical Approach

In order to apply both ETD and CCM to examine the dynamics of periodic and aperiodic EEG activity, we utilized the time series of epoched EEG data for delta, theta, alpha, beta, gamma and the aperiodic exponent (Figure 1B). This yielded 6 parameter time series of 896 time points (corresponding to each 2 second epoch, see Figure 1C). These time series were subject to both traditional delay-embedding and ETD (see Figures 1D and 1E) to determine the residual complexity for all parameters and all participants. We used repeated measures ANOVA to examine within participant effects of CONDITION (eyes open or eyes closed) and PARAMETER (aperiodic exponent or frequency band) as well as between participant effects of AGE (YA or OA) to examine effects of residual complexity.

We subsequently utilized EMCM to examine interactions between parameters and repeated measures ANOVA to examine within participant effects of CONDITION (eyes open or eyes closed) and PARAMETER (aperiodic exponent or frequency band) as well as between participant effects of AGE (young or old) to examine differences in dynamical interactions.

Lastly, we applied factor analysis to the resting-state New York State Cognition Questionnaire. This yielded a distribution of factors matching the results of the original study (Gorgolewski et al., 2014). We used multiple linear regression to examine relationships between attractor complexity and the form or content of resting-state thinking and corrected for multiple comparisons using the False Discovery Rate (FDR p<0.05).

## Results

### Aperiodic and Periodic EEG Parameters

Grand averaged EEG spectra for all participants is shown in Figure 1A. As expected, OA’s show a smaller aperiodic exponent, yielding a flatter aperiodic slope, which was most pronounced during the EO condition (AGE x CONDITION F(1,199)=25.55, p<0.00001, OAs: exponent_EC_=0.98; exponent_EO_=0.83 and YAs: exponent_EC_=1.15; exponent_EO_=1.10). For residual periodic power, we found a significant AGE X CONDITION interaction effect (F(5,995)=7.01, p=0.002). This was attributable to OAs showing less residual power in the theta and alpha bands (all corrected p’s <0.05), more residual power in the beta range (corrected p’s<0.00001) and more residual power in the gamma range in the EO condition only (corrected p=0.04). To examine the dynamics and geometric complexity of these parameters we next applied ETD embedding to yield spectral attractors.

### Spectral Attractor Complexity

Eigen-time-delay (ETD) embedding yields multidimensional manifolds that capture the trajectory of an individual EEG parameter’s dynamics over the course of a resting state scan. Although each parameter reveals idiosyncratic trajectories, there appeared to be recurrent dynamics traversed by each, suggestive of complex attractor dynamics (Figure 1E). Examining the ETD components for each parameter individually reveals quasiperiodic dynamics of increasing mean frequency and ranging from 0.01 to 0.1 Hz (Figure 2A, 2B, 2C). We also characterized the ETD components by their Maximal Lyapunov Exponent (MLE) to ascertain the stability of the underlying dynamics. We find a positive MLE across all components, suggestive of unstable dynamics, with the maximum exponent present at components 3 or 4 (Figure 2D). Thus, despite qualitatively idiosyncratic attractor trajectories, all EEG parameters for all conditions and groups show similarly unstable, quasiperiodic dynamics. Nonetheless, the percent variance explained by each ETD component was different for each parameter, condition and group. For YA participants, the first component of alpha during the EC condition captured the most variance (~15%). By contrast, for OA participants, the first component of the aperiodic parameter in the EC condition captured the most variance (~18%; See Figure 3). These differences in the distribution of explained variance resulted in differences in attractor complexity (number of retained components, see Methods) between parameters, conditions and groups.

**Figure 2.**
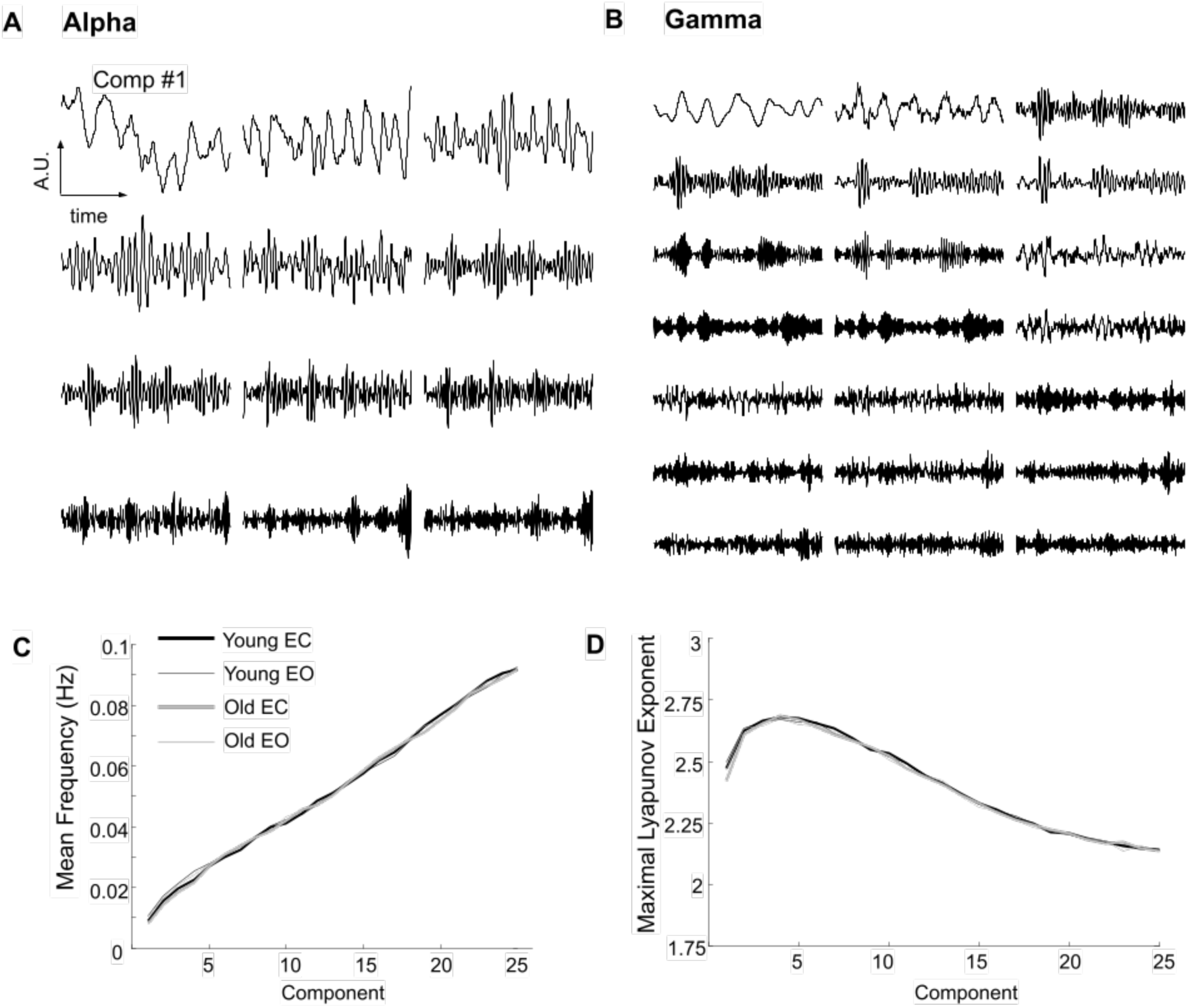
Eigen component dynamics. The first 12 principal components of alpha (A) and 21 components for gamma (B) for one participant. Components are ordered from left to right, ranked by the magnitude of the component eigenvalue. (C) Mean frequency of components (ordinate) plotted against the component number (abscissa) and averaged across all EEG parameters for YA (black) and OA (gray) participants during the EC (thick line) and EC (thin line) conditions. (D) Maximal Lyapunov Exponent (MLE, ordinate) plotted against the component number (abscissa) and averaged across all EEG parameters. Plotting convention is the same as in (C).

**Figure 3.**
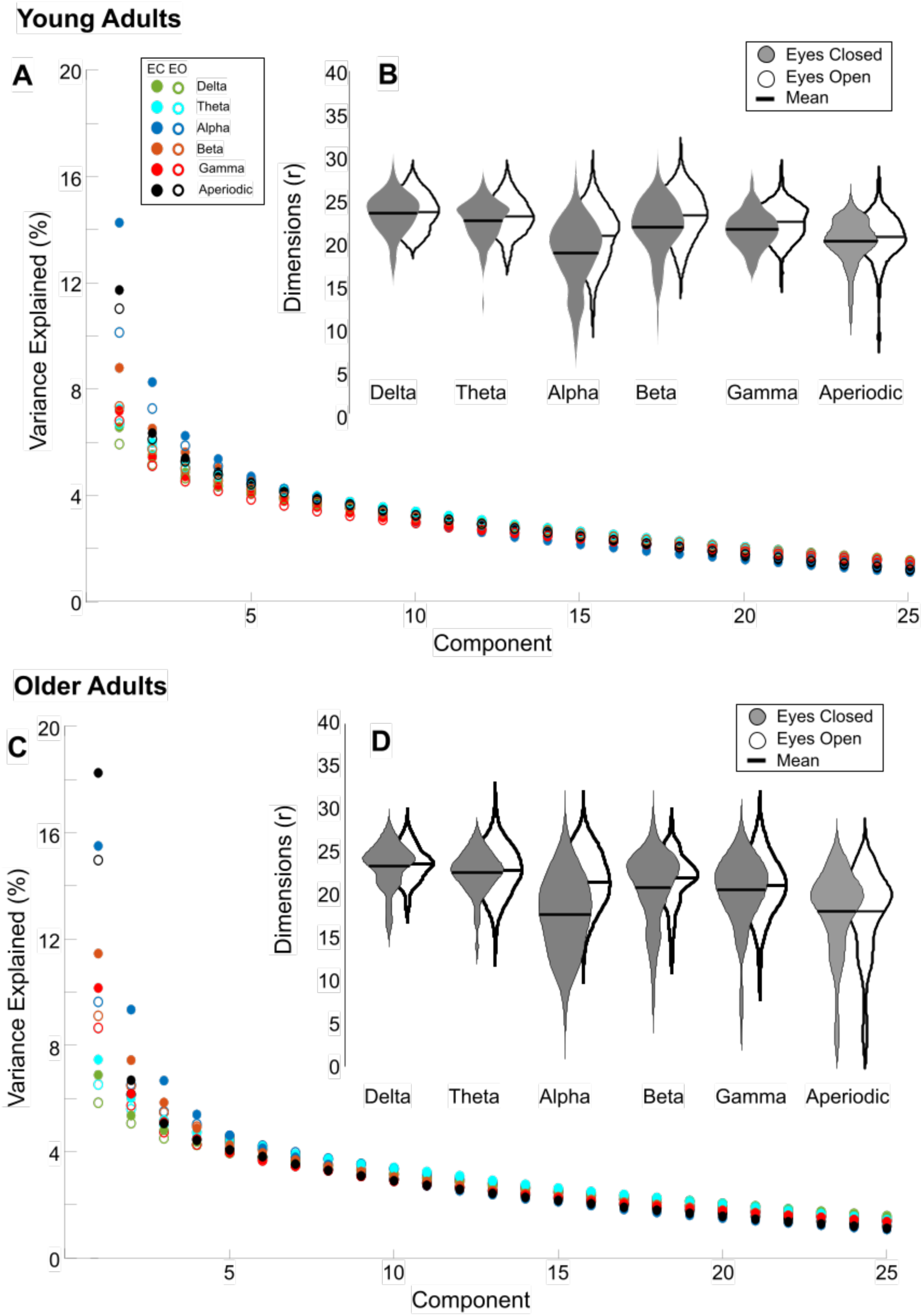
Parameter variance and geometric complexity. The average percentage of variance explained by each of the first 25 principal components (abscissa) across YA participants (A) and OA participants (C) for delta (green), theta (cyan), alpha (blue), beta (orange), gamma (red), and aperiodic (black), for eyes closed (filled circles) and eyes open (open circles). Violin plots display the distribution of retained principal components (geometric complexity) for each EEG parameter across YA (B) and OA participants (D) for eyes closed (gray) and eyes open conditions (white), with the mean complexity score indicated by the horizontal lines.

All participants show increased complexity during EO for periodic parameters (main effect of CONDITION, F(5,995)=63.6, p<0.0001) and the aperiodic exponent (main effect or CONDITION F(1, 199)=8.75, p<0.005). We found that the alpha attractors are the least complex relative to all other periodic parameters (main effect of PARAMETER, F(5,995)=17804, p<0.0001). Older adult participants show lower attractor complexity (main effect of AGE, F(1, 199)=5.63, p<0.05 and AGE X CONDITION X PARAMETER, F(5, 995)=2.47, p<0.05) in the beta and gamma attractors, during the EO condition (Tukey-Kramer corrected p<0.005) and gamma attractors in the EC condition (Tukey-Kramer corrected p<0.05). Older adults similarly show lower complexity in their aperiodic attractors across conditions (main effect of AGE, F(1, 199)=13.30, p<0.0005 without AGE x CONDITION effects).

In general, we found modest or no relationships between the magnitude of residual periodic power and attractor geometric complexity. When controlling for group and condition, greater alpha power modestly predicts less geometric complexity (partial r=−0.37, p<0.0005) with subtle, but significant differences in this association between young and old participants (rYoung=−0.27, pYoung<0.005 and rOld=−0.3, pOld<0.05; AGE*POWER Interaction (t-stat=5.07; p<0.0001). There was no significant relationship between delta, theta or beta power and corresponding geometric complexity (delta: r=0.02, p=0.73; theta:r=−0.05, p=0.34; beta:partial r=−0.06, p=0.20), and no significant differences in the relationship between groups, all ps >0.05). Only gamma attractors in young participants revealed significant associations between power and geometric complexity (YN: partial r=0.23, p=0.006; OA: partial r=−0.20,p=0.12, AGE*POWER Interaction (t-stat=1.99; p=0.047)). For aperiodic attractors, a larger aperiodic exponent was minimally correlated with greater complexity (partial r=0.16, p=0.001; without differences between groups, p=0.18). Thus the geometric complexity of spectral attractors is not simply a reflection of the mean power or mean aperiodic exponent, and reflects unique dynamical signatures. Next, we examined the local geometry of attractor dynamics and the extent to which these dynamics are shared across parameters.

### Geometric Cross-Parameter Coupling

To examine the magnitude of geometric cross-parameter coupling between spectral attractors we used Eigen Manifold Cross Mapping (EMCM, see methods). Low-dimensional, shared nonlinear dynamics between parameters on a participant-by-participant basis can be predicted using cross mapping methods (see Figure 4). EMCM (and CCM more generally) generates a predicted attractor based on a nearest neighbor approximation (see Methods and Figure 4) that is then correlated with the other parameter’s signal. Thus, EMCM provides a measurement of shared shape between dynamics, what we refer to as geometric cross-parameter coupling. We found that the highest correlations between predictor dynamics and the actual parameter dynamics were for the lower dimensions of the manifold, typically the first 15 dimensions which account for ~56-65% of the variance, (see Figure 5C). This finding suggests that a relatively low-dimensional set of core dynamics are shared across all EEG spectral parameters.

**Figure 4.**
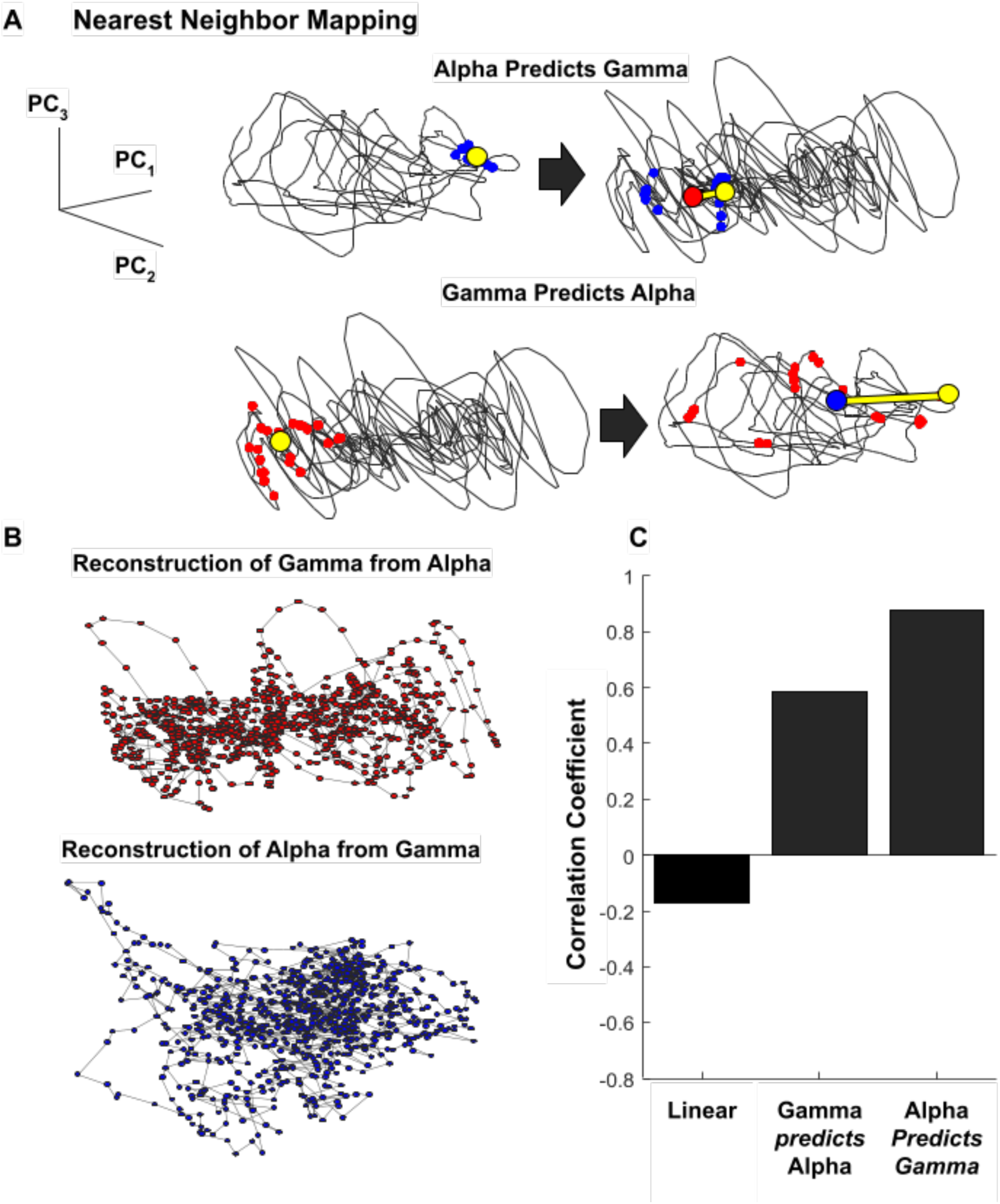
Geometric cross-parameter coupling is achieved by eigen manifold cross mapping. An example of geometric cross parameter coupling for a single participant (A) Top row, alpha predicts gamma: A random point (large yellow circle) on alpha’s attractor and its nearest neighbors (small blue circles) are mapped onto gamma’s attractor via the time-stamps of the neighbors on the alpha manifold (also designated by yellow and blue circles). The distance-weighted average of the mapped neighbors is designated by the large red circle and the total set of these averages are used to reconstruct a predicted gamma manifold by alpha (shown in B). The distance between the actual point on gamma (yellow circle) and the alpha’s prediction (red circle) is given by the thick yellow line; this distance is less than the corresponding prediction of alpha by gamma (see bottom row). Bottom row, gamma predicts alpha: utilizes the same convention as above. (C) Bar plot of the magnitude of Pearson correlation coefficient between alpha and gamma time-series (left), between the original alpha manifold and gamma’s manifold prediction of alpha (middle) and lastly, between the original gamma manifold and alpha’s manifold prediction of gamma (right).

**Figure 5.**
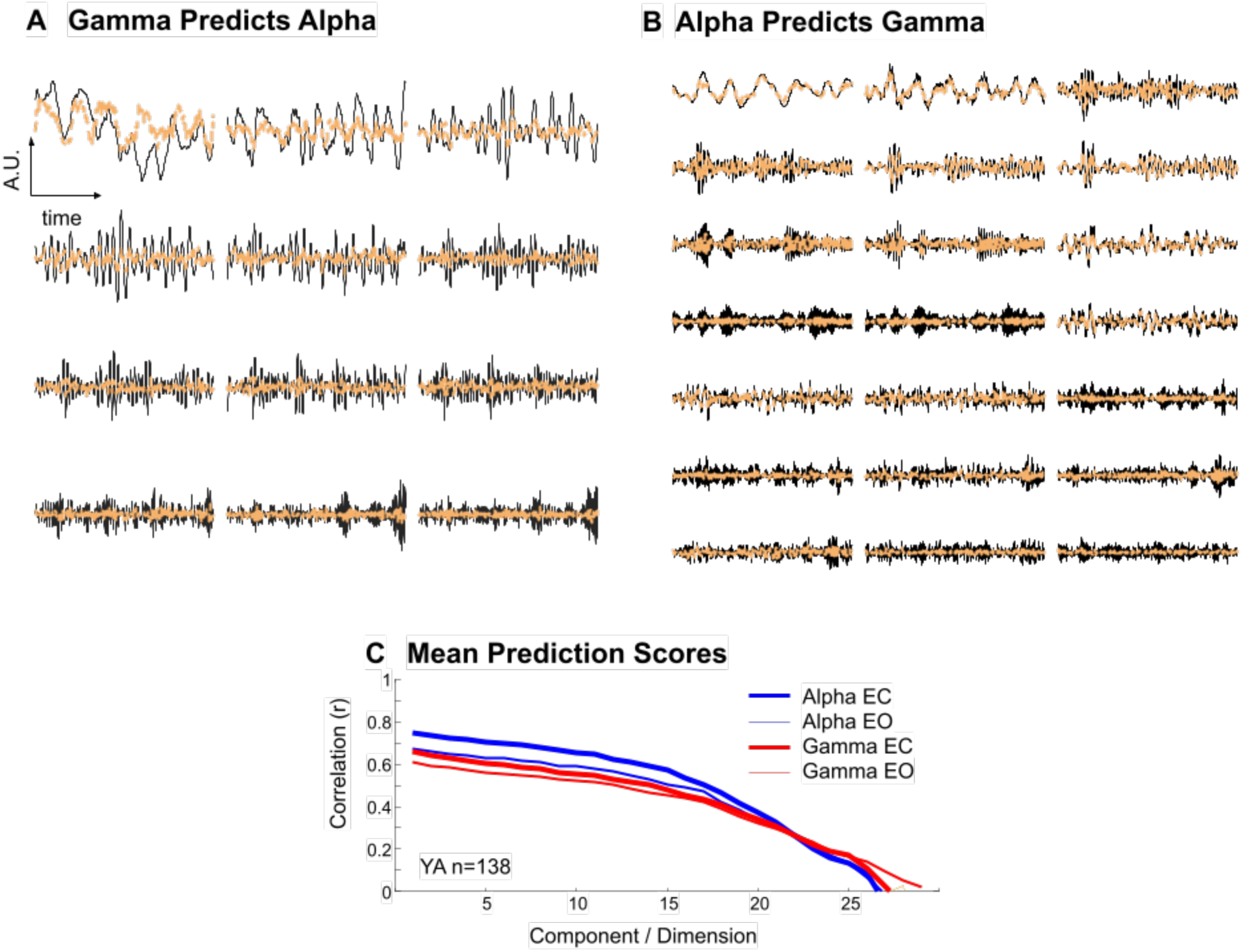
Low-dimensional gamma dynamics are predicted by alpha dynamics. (A) The dynamics of the first 14 principal components of alpha and their prediction by nearest neighbors of gamma’s manifold (dashed orange). (B) The dynamics of the first 21 principal components of gamma and their prediction by nearest neighbors of alpha’s manifold (dashed orange). (C) Average correlation coefficients between each component and its prediction for alpha (blue) and gamma (red) for EC (thick) and EO (thin).

To quantify the degree of geometric cross-parameter coupling, we report EMCM scores, which are the weighted average of all pairwise predictions between ETD dimensions. In general, we find moderate to high geometric coupling across all parameters, that is, most spectral attractors can predict the dynamics of the others (mean EMCM score of 0.56, range 0.26-0.99 across all participants and conditions). As described in the Methods and unlike linear correlation, (which yields a singular statistic; Pearson’s r for the pairwise relationship), EMCM yields two output statistics, one indicating the strength of prediction when signal *a* is the “predictor” and signal *b* “predicted” (*a* predicts *b*) and another when the signals are reversed and parameter *b* is the “predictor” and parameter a is “predictor” (*b* predicts *a*) Therefore, unlike a symmetrical linear connectivity matrix (Figure 6A, 6B), the top and bottom of the EMCM matrix are non-identical (Figure 6C, 6D). If we assume that each signal contains a dynamical signature of the core, an asymmetry might arise if one parameter is more “core-like” than the other.

**Figure 6.**
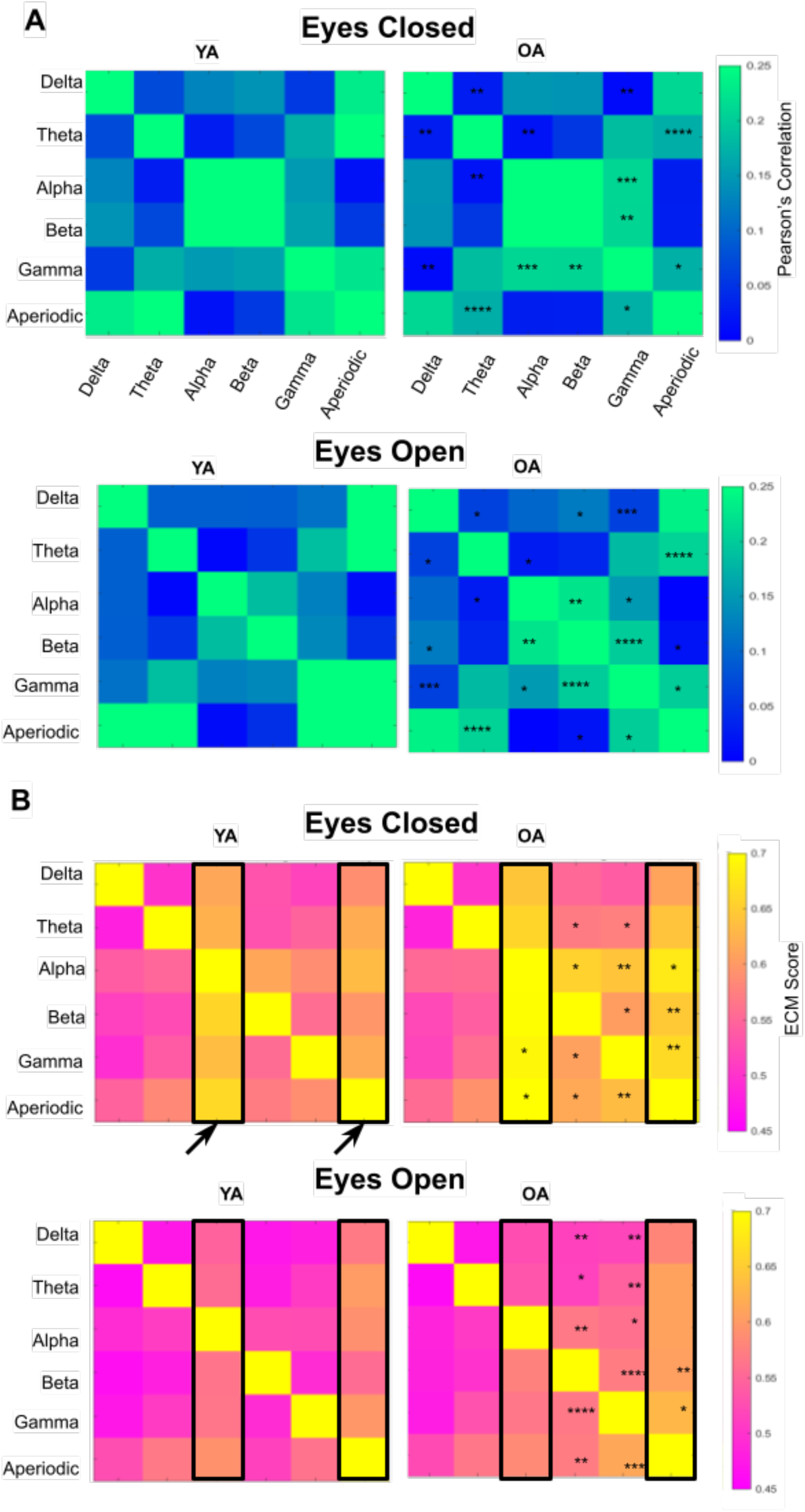
Linear correlation and geometric cross-parameter coupling by condition and group. (A) Linear correlation matrices for all parameters (delta, theta, alpha, beta, gamma, aperiodic), the magnitude of the absolute value of Pearson correlation coefficient is given by the colorbar for Young Adult (left) and Older Adult (right), EC (top) and EO (bottom). The asterisks signify the Tukey’s Honestly Significant Differences between YA and OA participants (*p< 0.05, ** p< 0.005, ***p< 0.0005, **** p< 0.00005) (B) EMCM matrices for all parameters (delta, theta, alpha, beta, gamma, aperiodic), the magnitude of the EMCM score is given by the colorbar for Young Adult (left) and Older Adult (right), EC (top) and EO (bottom). Significant differences are indicated by asterisks, as in (B). Arrows and boxes denote core-like, shared dynamics.

Across conditions, and in YA and OA alike, alpha predicts all other spectral attractors the best (main effect of PARAMETER, F(1,5771)=124.27, p<0.0001; Figure 6C & D, vertical yellow bands, Tukey’s HSD for all alpha prediction p’s < 0.0001). In geometric terms, this means that the alpha attractor contains signatures that are the most shared among all other attractors (i.e., alpha has the most generic dynamical signatures). The aperiodic exponent also exhibits this pattern, whereby most other parameters show greater EMCM scores when they are predicted by the aperiodic parameter (Tukey’s HSD for all aperiodic prediction p’s (except alpha) < 0.0001). Older adults show greater EMCM scores across conditions (main effect of AGE, F(1,5771)=6.61, p=0.010). This is most pronounced when the higher frequency attractors and the aperiodic exponent are predictors (AGE x CONDITION x PARAMETER F(1,5771)=2.63, p<0.0001; significance of follow-up tests using Tukey’s HSD are shown in Figure 6).

In general, we find that EEG attractors with reduced dynamical complexity are better predictors, that is, their dynamics are most similar to the set of low-dimensional, core-like dynamics. This was true across all participants and conditions. Alpha attractor complexity inversely correlates with EMCM scores across parameters while controlling for age and condition (all partial r’s <-0.86 and all p’s <0.0001, and no AGE x CONDITION effects). In contrast, the geometric complexity of the other parameters was only modestly associated with lower EMCM scores. (delta complexity r=−0.46, theta, r=−0.37, beta, r=−0.56, gamma r=−0.45, exponent=−0.42, all p’s<0.0001). Similar to alpha, aperiodic attractor complexity inversely correlates with EMCM scores (all partial r’s <-0.79 and all p’s <0.0001, no AGE x CONDITION effects) and where dynamical complexity of the other bands only modestly predicts lower EMCM scores (delta: r= −0.50, theta : r=−0.54, alpha: r=−0.52, beta: r=−0.60, gamma complexity : r=−0.58, all p’s<0.0001). These correlations are entirely eliminated if the complexity of the opposite condition attractor is used (e.g. the complexity of the EO alpha attractor cannot predict the EMCM scores of EC delta, theta, beta and gamma). This suggests that the correlation between complexity and prediction scores depends on the unique condition-dependent geometric structure of the spectral attractors. That this correlation was most robust for alpha and aperiodic, suggesting that these two parameters contain the core-dynamics of the EEG spectrum at rest.

### Correlations with Resting State Thoughts

We hypothesized that the geometric complexity of EEG attractors might be associated with participants’ mental state at rest. To explore this possibility, we utilized responses to the New York Cognition Questionnaire collected from a fMRI session the day before the EEG session. We reasoned that the form and content of mental activity at rest would be habitual, automatic and recurring (Colvin et al., 2021; Hertel, 2004; Watkins & Nolen-Hoeksema, 2014) and well captured by the attractor paradigm (Adams et al., 2022). Thus, although we do not have access to the form and/or content of the participants mental state during the EEG recording, we hypothesized that there might be some similarity between the resting state mental state during EEG and fMRI sessions. Following Gorgolewski et al. we used factor analysis to identify dimensions of content and form, yielding similar factor loading matrices (Gorgolewski et al., 2014).

OA participants show lower factor scores related to thoughts about the past or future (t=3.30, p=0.001, t=5.35; p<0.001) as well as lower factor scores about positive thoughts (t=2.96, p=0.003), but not negative thoughts (t=1.23, p=0.21). OAs show higher factor scores related to thoughts about friends (t=−3.32, p=0.001). Regarding form, older adults show lower factor scores related to imagery (t=5.04, p<0.0001) and vagueness (t=5.58, p<0.0001) but not verbal thoughts (t=0.96, p=0.34). Given the exploratory nature of this analysis, we included the complexity of all EEG parameters as predictors of form and content in repeated measures, multiple linear regression analysis that included age and condition as covariates. After controlling for multiple comparisons, only the complexity of gamma band attractors was associated with factor scores for negative thoughts (p_fdr_=0.016, partial r_young_=0.21, r_old_=0.17) without differences in the magnitude of this correlation between groups (p_unc_=0.11). With the exception of aperiodic attractor complexity, there were no interaction effects between parameter complexity and age in predicting mental state. For OAs (but not YAs) the complexity of the aperiodic attractor correlates with the factor scores for positive thought content (partial r_young_=−0.12, r_old_=0.25, AGE x COMPLEXITY p_fdr_=0.017) and visual imagery (partial r_young_=−0.11, r_old_=0.31, AGE x COMPLEXITY p_fdr_=0.017).

## Discussion

In summary, we have implemented eigen-time-delay embedding to characterize the geometric complexity of EEG spectral dynamics. This approach examines an EEG parameter’s signal in state-space and uses PCA to determine the minimum number of components (dimensions) necessary to capture those dynamics. The output of this approach yields a lower-dimensional attractor that describes the “shape” of an EEG parameter’s dynamics. The local geometry of this shape is characterized by the Lyapunov exponent (instability) and the extent to which local neighborhoods can predict the location of neighborhoods on another parameter’s manifold, what we refer to as geometric cross-parameter coupling. These analyses reveal that the two hallmarks of ongoing activity in the resting EEG spectra, alpha power and the aperiodic exponent, contain geometric signatures of all other parameters. We refer to these parameters as “core,” consistent with the view that specific cognitive operations emerge from ongoing activity in a dynamically unstable dominant assembly (Le Van Quyen, 2003; Varela, 1995). All parameters contain core-like activity that is low-dimensional, unstable, and suggestive of attractor dynamics. However, non-core parameters (delta, theta, beta and gamma) are characterized by higher-dimensional manifolds, with the geometric complexity of the gamma band reduced in older adults and associated with resting-state mental content. This distinction between core and non-core dynamics suggests that EEG spectral dynamics reflect a spectrum from more integrative (alpha/aperiodic) to more differentiated (gamma) and that geometric approaches can be employed to characterize the degree of integration and differentiation (Tononi & Edelman, 1998). Next, we interpret these findings in the context of dynamical systems theory, geometric interpretations of neural data and theories of cognitive aging.

### Geometric Attractor Complexity

We have developed a measure of geometric complexity that distinguishes between periodic and aperiodic EEG dynamics, depends on the state of the visual system (greater complexity in EO vs EC) and identifies a loss of complexity in aging. Our geometric assessment of complexity is based on the embedding dimension in state space, extending prior non-human primate work in ECoG that found greater geometric complexity in the awake compared to the anesthetized state (Tajima et al., 2015). This approach is broadly consistent with the perspective that brain dynamics operate in a high-dimensional space (Tozzi, 2019), and where neurophysiological recordings reflect low-dimensional observations of multi-dimensional dynamics (Elsayed & Cunningham, 2017; Gallego et al., 2017; Nogueira et al., 2023; Shenoy et al., 2011). Geometric frameworks have been applied to the readout of multi-unit neuronal recordings with potential for decoding population activity (Azeredo da Silveira & Rieke, 2021; Chung & Abbott, 2021; Jazayeri & Afraz, 2017; Jazayeri & Ostojic, 2021). In multi-unit neuronal studies, 12-dimensional manifolds capture upwards of 73% of the task-based variance (Gallego et al., 2017, 2018). Our assessment of EEG data during the resting state finds that the first 15 components capture ~64-71% of the variance for alpha and aperiodic and ~55-62% of the variance for non-core bands. More work is needed to bridge the perspective of population activity with EEG dynamics (P. Gao & Ganguli, 2015), but these findings suggest that population activity, ECoG and EEG manifolds show similar complexity (10-15 dimensions), despite widely varying tasks, conditions and preparations. One possibility suggested by (Pinotsis & Miller, 2022) is that the low dimensionality of EEG reflects stable electric fields that funnel population activity along lower dimensions, a concept that had previously been proposed by Freeman via carrier waves (Freeman, 2003).

Although there is a long history of applying state-space embedding methods to EEG analysis (Freeman, 1987; Tsuda, 2001), geometric analyses of manifolds have only recently been applied, and generally to the dynamics of the whole EEG or ECoG spectrum (Akbari et al., 2021; Kwessi & Edwards, 2021; Varley et al., 2021). However, EEG dynamics reflect a mixture of periodic signals across a range of frequency bands, in addition to contributions from aperiodic components, sometimes referred to as scale-free since this activity lacks a dominant temporal frequency. Therefore, rather than representing a singular statistic for the entire EEG, complexity scores may vary from parameter to parameter, which informed the approach we took here. Critically, we find that alpha and aperiodic parameters are the least complex whereas low and high frequency activity, (on either side of the alpha peak) show greater geometric complexity. This is consistent with a widely employed heuristic that a shift toward higher frequency activity is indicative of brain activation (Kilner et al., 2005), as gamma band power has been studied extensively in relation to cognitive tasks (Fries et al., 2008; Herrmann et al., 2010; Melloni et al., 2007) and as a putative marker of E/I balance in resting-state studies (White & Siegel, 2016). Thus, our finding of increased complexity in the beta and gamma bands is consistent with these bands’ putative role in more complex cognitive tasks. Further, that gamma complexity was the only parameter associated with thought content during the resting state task, is consistent with evidence linking gamma band activity to ruminative thinking (negative, recurring patterns of thought) (Wang et al., 2022). That is, if the resting state represents a diverse sampling of states of mind (Diaz et al., 2013; Gorgolewski et al., 2014), then we speculate that participants with greater gamma complexity may have been more engaged with ruminative thinking. This hypothesis is supported by a shift toward lower frequency activity during states of mindfulness, a period of reduced ruminative mental activity (Lomas et al., 2015).

### Geometric Cross-Parameter Coupling

Our use of geometric complexity enables a novel analysis of cross-parameter coupling based on the geometry of local dynamics for each EEG parameter. Whereas linear correlation approaches find modest relationships between nearby EEG parameters (most notably alpha and beta), the geometric approach we utilized finds that alpha and aperiodic signals show the most robust coupling between other EEG parameters. Cross-parameter (typically cross-frequency coupling) is a well established analytic approach to task-based brain-cognition-behavior relationships (Aru et al., 2015; Canolty & Knight, 2010; Hyafil et al., 2015; Palva & Palva, 2018). A related concept of intrinsic coupling has been advanced to examine resting or ongoing brain activity that considers either band-limited phase coupling or coupling in the envelope (amplitude) of signal fluctuations (Engel et al., 2013). Our application of geometric cross-parameter coupling adopts the latter approach, embedding the signal envelope of EEG parameters into state-space. This enables us to examine potential interactions across a range of periodic and aperiodic signals. More importantly, it allows us to examine nonlinear and directed interactions between neural signals from cross-mapping methods (Tajima et al., 2015).

Convergent cross-mapping (CCM) was developed by Sugihara and colleagues to infer directed-causal interactions between highly nonlinear signals (Sugihara et al., 2012; Sugihara & Ye, 2009; Ye et al., 2015). Subsequently, CCM has been applied to ECoG recordings in non-human primates (Tajima et al., 2015), multielectrode recordings in rats (Cocina et al., 2021) and to a limited extent in human EEG (Fonseca et al., 2018; Lainscsek et al., 2019; McBride et al., 2015). Similar to (Tajima et al., 2015), we find a high degree of bi-directional interaction between neural signals that would be otherwise missed from purely linear approaches. Alpha and aperiodic parameters stand out in their ability to predict the nonlinear dynamics of all other frequency bands. In the language of Sugihara causality, this means that all other frequency bands have imprinted their dynamical signatures on these core bands (i.e. caused their dynamics). Core bands are the least complex and the most dynamically generic and accommodating, suggesting that they have been involved in more processes. This perspective is broadly consistent with contributions of alpha and aperiodic signals to general arousal (Colombo et al., 2023; Jacob et al., 2021; Lendner et al., 2020; Ota et al., 1996) and alpha’s role as a brain-wide inhibitory signal across task and resting states (Jensen & Mazaheri, 2010).

Since all EEG parameters share geometric dynamics with alpha and aperiodic signals, this motivates our interpretation that these dynamics represent a geometric core of resting-brain activity. Accordingly, the EEG spectra can be envisioned as “sew-saw,” with the fulcrum at the alpha band (Garcia-Rill et al., 2016) and all rhythmic activity emerging from the beam of aperiodic activity (Figure 7A). All other frequency-based parameters differentiate out of the more integrative alpha/aperiodic dynamics to yield higher complexity signals based on the nature or content of the ongoing mental state (Figure 7B, 7C). A dynamic core hypothesis originates with the work of Varela (Le Van Quyen, 2003; Varela, 1995), was extended by Tononi and Edelman (Tononi & Edelman, 1998) and has largely been explored theoretically (Deacon, 2011). Dynamic core models have also been linked to thalamocortical inhibitory dynamics (Min et al., 2020; Ward, 2011), which is highly supportive of our finding alpha as the dominant signal of the dynamic core. Our results provide electrophysiologic evidence for the dynamic core hypothesis from the shared dynamics of cross-parameter interactions during the resting state. Our approach reveals that core dynamics are not merely indexed by linear phase or amplitude synchronization, but by nonlinear, geometric features in ongoing signal dynamics. Critically, both the alpha and aperiodic signals are thought to be expressions of arousal, E/I balance and/or global inhibition (R. Gao et al., 2017; Jensen & Mazaheri, 2010; Klimesch et al., 2007; J. M. Rhodes, 1969). Therefore, we hypothesize that core dynamics are more than gain controls for cortical activity (as might be suggested by the E/I balance model) but identify diverse geometric shapes that reflect the corresponding diversity of fluctuating, multidimensional states of arousal and awareness (Munn et al., 2021; Thayer, 1978; Trofimova & Robbins, 2016). Future work will be needed to examine this hypothesis through task-based experimental manipulation.

**Figure 7.**
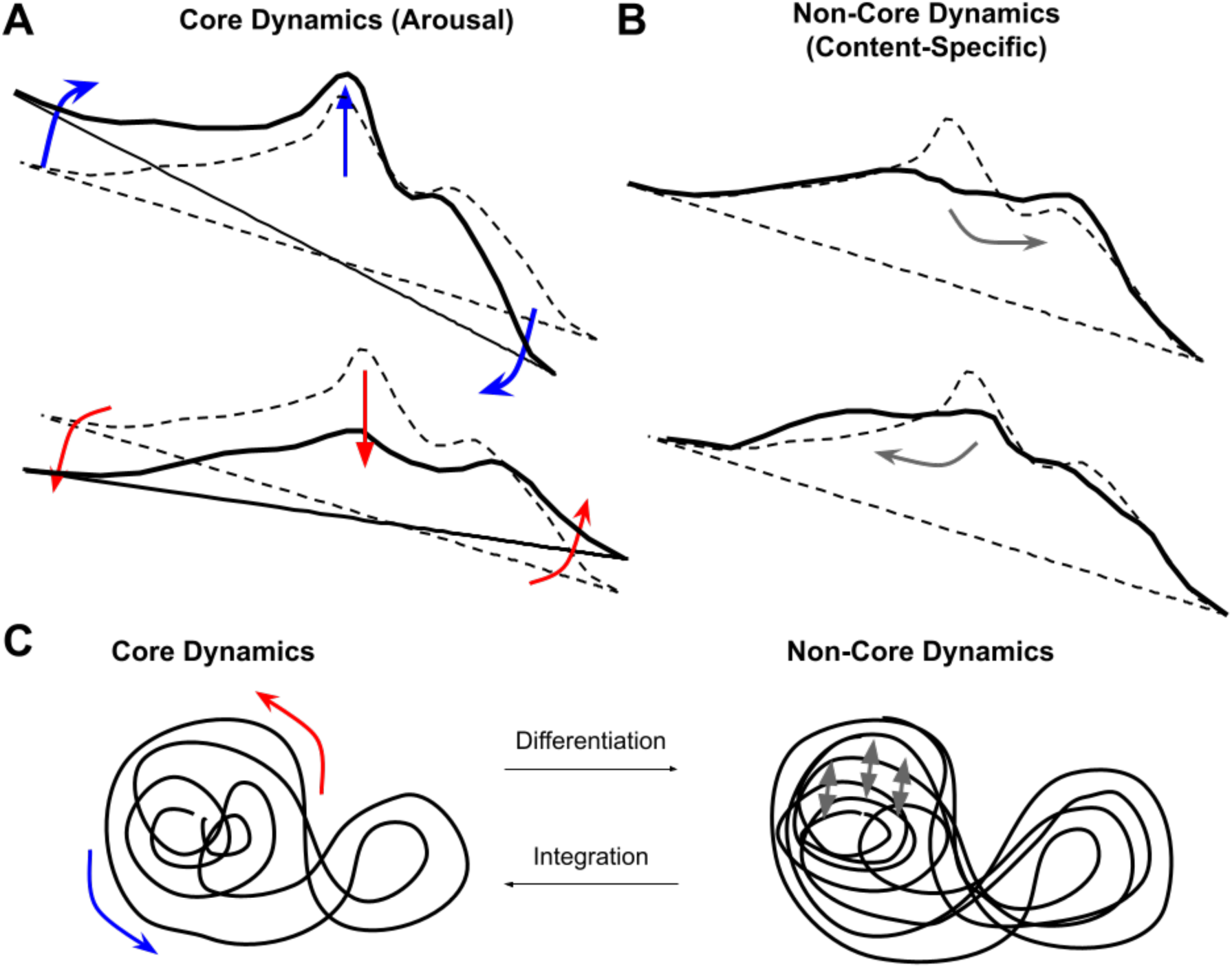
Schematic of spectral features underlying the dynamics of the geometric core. (A) Moment-by-moment fluctuations in arousal underlie core dynamics via changes in the aperiodic slope and the magnitude of alpha power. Top: Decreasing arousal and global inhibition increase the aperiodic slope and alpha power (blue arrows). Bottom: Increasing arousal and global excitation decrease the aperiodic slope and alpha power (red arrows). (B) Moment-by-moment processing of content-specific, mental-state dynamics arises via higher-complexity, non-core dynamics. This can occur via shifts toward increased high-frequency power (top) or lower frequency power (bottom). (C) Core-dynamics are lower complexity, and therefore more integrative (left). Non-core dynamics differentiate out of core dynamics, yielding more complex manifolds (right).

### Loss of Complexity in Aging

Similar to others, we have found a smaller spectral exponent (shallower slope) in older adults (Donoghue, Haller, et al., 2020) and a reduction in alpha power (Breslau et al., 1989; Roubicek, 1977; Vysata et al., 2012), further supporting the importance of decomposing periodic and aperiodic components for studies of EEG activity in aging (Tröndle et al., 2023). Despite these mean differences in alpha power, we find no differences in alpha complexity between YA and OA. By contrast, we find no differences in gamma or beta periodic power, however, OAs show reduced complexity of gamma and beta and aperiodic complexity. This suggests that the complexity of core dynamics remains largely intact in late life, and in particular, alpha remains a robust predictor of other frequency bands. Nonetheless, the loss of complexity in beta, gamma and aperiodic parameters occurs concomitant with an increase in geometric coupling scores between these parameters. We interpret this finding as a loss of high-frequency signal differentiation, that is, the aging brain is more dynamically homogenous, less complex but more causally efficacious.

This interpretation is consistent with two well established and related theories of brain aging, including the loss of complexity in aging and the dedifferentiation hypotheses. The loss of complexity in aging hypothesis proposes that a wide range of physiologic signals show reduction in variability, richness of connectivity and the degree of hierarchical organization (Lipsitz & Goldberger, 1992; Ma et al., 2021; Vaillancourt & Newell, 2002). In the aging brain, loss of complexity might result from gray matter and synapse loss, leading to decreased long-range connectivity (Huttenlocher, 1979; Masliah et al., 1993; Scheff & Price, 2003). This process may be related to reduced specificity of regional activation or broadening of tuning curves, generally interpreted as a dedifferentiation in aging (Heuninckx et al., 2008; Park et al., 2004; Rakesh et al., 2020; Sleimen-Malkoun et al., 2014). Our finding that OA participants show reduced geometric complexity along with greater geometric coupling, is supportive of the signal dedifferentiation hypothesis. This mirrors prior findings of exaggerated alpha-gamma synchronization between nodes of the default mode network in older adults during the resting-state, associated with impaired working memory performance and interpreted as being “stuck” in default mode (Pinal et al., 2015). Enhanced synchronization and cross-parameter coupling could reflect compensatory mechanisms to sustain neural dynamics in the face of reduced signal power or complexity (Ansado et al., 2012; Jacob & Duffy, 2014; Meunier et al., 2014; Morcom & Henson, 2018; Zahodne & Reuter-Lorenz, 2019). Longitudinal studies are needed to determine whether increased synchronization precedes the loss of complexity or results from a compensatory process.

### Limitations and Future Directions

This study has taken a novel, geometric approach to analyze the dynamics and cross-parameter coupling of EEG spectral data. We have made use of a publicly available dataset to demonstrate the applicability of this approach to non-clinical, human populations and normal aging. A primary limitation of this study is that we have only examined resting-state data. Our approach does not preclude application to task-based experimental design, however, sufficiently long task blocks would be required to capture the EEG spectral manifold and cross-mapping interactions (Maher & Hernandez, 2015). Nonetheless, that we find differences between EC and EO conditions as well as correlations with the content or resting-state thoughts, suggests state-dependence of both geometric complexity and cross-parameter coupling. Our application of the dynamic core concept complements fMRI-derived (BOLD) dynamics that are consistent across diverse cognitive tasks (Shine et al., 2019), by examining dynamics that are consistent across EEG parameters. For task-based applications, one could modify the pipeline we have proposed here to identify principal dimensions of interest based on experimental or task variables, or to compare against linear predictions (Brunton et al., 2017).

Our study is also limited by the fact that the resting-state cognition questionnaire was conducted the day before the EEG session, and during an fMRI session. Differences between EEG and fMRI experimental conditions notwithstanding, participants might have had different thought patterns and content during the two sessions. Our analysis was exploratory, and so this finding must be interpreted with caution. Nonetheless, it is reasonable to consider that similar thought patterns may persist from day to day (Colvin et al., 2021; Hertel, 2004; Watkins & Nolen-Hoeksema, 2014). Future work will be needed to examine if such habits of mind can be detected from the geometric shape of brain activity.

Lastly, we have not conducted an exhaustive comparison with information theoretic approaches that evaluate complexity based on statistical redundancies in the data (e.g. Kolmogorov, Lempel-Ziv or multiscale entropy) (Cafaro & Ali, 2021; Wang, 2012)). Information-theoretic approaches have not distinguished aperiodic and periodic contributions to the EEG spectra (Al-Nuaimi et al., 2018; González et al., 2022; Pappalettera et al., 2023; Sun et al., 2020), thus future work might identify differences between information-theoretic and geometric approaches to EEG spectral data. We utilized a geometric approach because of increasing evidence that variability in brain activity can be characterized by dynamic evolution in manifold structure. These approaches might herald a “geometric turn” to examine neural activity not as a statistical code, but according to resemblances among iconic shapes that characterize a diversity of trait and state-based dimensions in mental experience (Deacon, 2022; Williams & Colling, 2018).

## Acknowledgements

This work was supported by grants from the Department of Veterans Affairs VA (IK1CX002089 and IK2CX002457 to MSJ). We gratefully acknowledge the input from our colleagues, who provided valuable comments on an earlier version of this manuscript: Terrence Deacon, Matthew Rogers, Charles Duffy, Adam Jacob, Mani Hamidi and Judith Ford.

